# Autophagy is induced during plant grafting for wound healing

**DOI:** 10.1101/2020.02.14.949453

**Authors:** Ken-ichi Kurotani, Ryo Tabata, Yaichi Kawakatsu, Ryohei Sugita, Koji Okayasu, Keitaro Tanoi, Michitaka Notaguchi

**Affiliations:** Bioscience and Biotechnology Center, Nagoya University, Furo-cho, Chikusa-ku, Nagoya 464-8601, JAPAN; Graduate School of Bioagricultural Sciences, Nagoya University, Furo-cho, Chikusa-ku, Nagoya 464-8601, JAPAN; Isotope Facility for Agricultural Education and Research, Graduate School of Agricultural and Life Sciences, The University of Tokyo, Yayoi, Bunkyo-ku, Tokyo 113-8657, JAPAN; Institute of Transformative Bio-Molecules, Nagoya University, Furo-cho, Chikusa-ku, Nagoya 464-8601, JAPAN

**Author notes:** These authors contributed equally to this work (Ken-ichi Kurotani, Ryo Tabata, Yaichi Kawakatsu). Author for correspondence: Michitaka Notaguchi.

**Keywords:** Arabidopsis, Autophagy, *atg2*, ATG5, ATG8, Grafting, *Nicotiana benthamiana*, wound healing

## Abstract

- Grafting is an important technique in agriculture to obtain several good traits such as high disease tolerance and high yield by exchanging root system. However, the underlined cellular processes to compensate the wound damage and repair tissues were largely unknown.
- We analyzed two graft combinations: *Nicotiana benthamiana* (*Nb*) homograft as a compatible, wound repairing model and *Nb* heterograft with *Arabidopsis thaliana* (*At*) as an incompatible and more stressful model, which we recently identified as an exceptional maintainable interfamily grafting.
- In both graft combinations, nutrient loss was observed in gene expression after grafting, where the level of nutrient loss was more sever in heterografts. Transmission electron micrographs of *Nb*/*At* heterografts suggested that microautophagy was induced in cells near the graft boundary. In *At* seedling micrografting, the fluorescence of autophagy protein marker GFP-ATG8 was highly observed at graft junction especially in cambial region. In *At atg2* mutant homografts, growth after grafting decreased compared with wild-type homografts. Moreover, when *NbATG5* knocked-down *Nb* scion was grafted to *At* stock, the successful rate of grafting was significantly decreased.
- Altogether, these results suggest that component of autophagy is induced during grafting and has a role in wound healing.

## Introduction

Grafting is an important method in agriculture to propagate heterogenic but attractive tree species and/or to take benefit of rootstock traits such as disease tolerance and stress tolerance to the unfavorable soils (Davies *et al*., 2018). Grafting is essentially dependent on the native ability of wound repairing in plants (Wang, 2011; Goldschmidt, 2014). The molecular mechanism of wound repairing on the hypocotyl or the stem was examined in cucumber, tomato and Arabidopsis (Asahina & Satoh, 2015). Asahina *et al*. (2002) suggested that gibberellin (GA) is required for cell division in the cortex of wounded cucumber and tomato hypocotyls. Further physiological and molecular analyses suggested that auxin, ethylene and jasmonic acid contribute to the wound healing process of the Arabidopsis stem through the action of a couple of transcription factors functioning in phytohormone signaling pathway (Asahina *et al*., 2011; Pitaksaringkarn *et al*., 2014). These studies indicated that endogenous phytohormones levels are important for tissue reunion after wounding. Similar idea was also obtained through the studies of grafting. Recent study using Arabidopsis as a model showed that auxin and a subset of auxin response genes have important role for graft establishment (Melnyk *et al.*, 2015; Matsuoka *et al.*, 2016). Not only phytohormone action, many of the genes related to metabolism, the oxidative stress response, transport, and defense are involved in graft healing process. Transcriptomic analyses conducted on the grafts of fruit trees (Chen *et al.*, 2017; Assunção *et al.*, 2019) and the seedling grafts of Arabidopsis (Melnyk *et al.*, 2018) and tomato (Wang *et al*., 2019; Xie *et al.*, 2019) identified that graft incision induced drastic transcriptomic changes including upregulation of the wound response genes after grafting.

On the other hand, structural studies of grafting processes demonstrated that grafting is achieved through wound response, callus bridge formation, adhesion, differentiation of wound-repairing vascular tissues and production of secondary xylem and phloem in the callus bridge (Davies *et al*., 2018). Ultrastructural analyses on the graft boundary region showed that a necrotic layer of one or two damaged cells initially lays between the stock and scion and the layer was fragmented by 2 to 3 days after grafting as the callus proliferation continued (Moore & Walker, 1981). Newly proliferated callus from the peripheral tissues produced fibrillary network and bead-like structures on the cell surfaces and the extracellular material subsequently sealed the graft union (Jeffree & Yeoman, 1983; Sala *et al*., 2019). Plasmodesmata were newly formed in the cell wall region at graft boundary, especially the site where vascular tissues were well aligned among the stock and scion (Kollmann *et al*., 1985). Thus, a sequence of cell adhesion processes during wound healing has been well described.

In this study, we investigated the events inside of the cells during grafting by transmission electron microscopy and found the typical structures observed in the autophagic plant cells. In nature, plants are often affected by internal and external stresses. Lack of essential nutrient elements which frequently occurs threatens plant survival, e.g. carbon deficiency caused by the limit of photosynthetic efficiency (Bassham *et al*., 2006), nitrogen or phosphate deficiency due to low concentrations in the soil (Maathuis, 2009). Autophagy is a key process as a function to counter these deadly stresses in plants (Bassham *et al*., 2006). In eukaryotic cells, autophagy is a universal mechanism for discarding damaged or harmful components and for recycling cellular materials during development (Farré & Subramani, 2016; Li *et al*., 2012; Liu & Bassham, 2012). The function of autophagy has been revealed by morphological observations with the electron microscope. In cells where autophagy appears to be progressing, bulk sequestration of cytoplasmic fragments and subsequent digestion in the lytic vacuoles were observed. Three distinct types of autophagy have been reported in plants: micro-autophagy, macro-autophagy and mega-autophagy (Bassham *et al*., 2006; van Doorn *et al*., 2013). The details of micro-autophagy are not well understood, but cytoplasmic components that come in contact with the surface of vacuole are directly engulfed by invagination of tonoplast and taken into the vacuole as vesicles called autophagic bodies (Bassham *et al*., 2006). On the other hand, macro-autophagy in plants has been studied more detail and core machinery has been described. Macro-autophagy forms autophagosomes, the double-membrane bound structure that enwrap cytoplasmic components such as endoplasmic reticulum, Golgi stacks, mitochondria, plastid and peroxisomes. Then these vesicles are transported to vacuole interior by fusing the autophagosomal outer membrane to vacuole surface, followed by digestion in vacuole (Liu & Bassham, 2012). Mega-autophagy is involved in the cell destroy machinery of developmental programmed cell death, such as xylem formation in plants (van Doorn *et al*., 2013; Kwon *et al*., 2010).

The molecular mechanism of autophagy is best understood in macro-autophagy. Macro-autophagy depends on approximately 40 core Autophagy-related (ATG) genes that are highly conserved from yeast to mammals and plants. In plants, many ATG transcript levels are increased by the senescence and nutritional starvation. Among them, ATG3, ATG4, ATG5, ATG7, ATG8, ATG10, ATG12, ATG16 constitute two ubiquitin-like conjugation systems essential for autophagosome formation (Yoshimoto, 2012; Soto-Burgos *et al*., 2018). ATG8 forms a conjugate with phosphatidylethanolamine and associate to membrane tissue. This conjugate abundance is often used as an indicator of autophagy activity, and GFP-tagged ATG8 is used as a tool to monitor the autophagy mechanism (Yoshimoto, 2012). ATG2 forms a complex with ATG9 and ATG18 which plays an important role in lipid transport from ER to phagophore (Zhuang *et al*., 2017; Gómez-Sánchez *et al*., 2018). On the other hand, little is known about the molecular mechanism of micro-autophagy, but the essential roles in plant life has been unveiled. Micro-autophagy functions in the formation of cytoplasmic anthocyanin deposition to the vacuoles in epidermal cells and chlorophyll degradation during leaf senescence in both core ATG machinery-dependent and -independent manners (Krick *et al*., 2008; Chanoca *et al*., 2015; Izumi *et al*., 2019; Nakamura & Izumi, 2019). When plants were subjected to strong external stresses, degradation of chloroplasts in vacuole were sometimes observed. In such cases, it is known that stroma of chloroplast is surrounded by a membrane to form small vesicles referred to as Rubisco-containing bodies (RCBs), in other hand, whole chloroplasts including stroma and thylakoids are also subjected to micro-autophagy termed chlorophagy. In plants subjected to photodamage, damaged whole chloroplasts were observed to associate with structures containing ATG8 and were engulfed by tonoplast and to be transported to the vacuolar lumen. In mutants of ATG5 and ATG7, these phenomena were not observed, indicating chlorophagy is dependent on the core ATG machinery (Nakamura *et al*., 2018; Izumi *et al*., 2017; Izumi *et al*., 2019). Failure of autophagy activation in mutants showed short life phenotype because of early senescence and less fitness to environmental light conditions (Bassham *et al*., 2006). Thus, the importance of autophagy activation in plant growth and development has been clearly demonstrated and further insights on the function of autophagy in plants will facilitate plant biology.

In this study, we investigated how do plants compensate graft wound situation and repair the tissues for their survival. Genetic distances of grafted plant species strongly affect the efficiency of repairing at the grafted tissues and generally most successful grafting has been conducted in same genus/family plants where wound healing process is accomplished within short period, newly vascular formation starts within several days (Davies *et al*., 2018; Wang, 2011; Goldschmidt, 2014). However, we previously identified *Nicotiana benthamiana* (*Nb*) and *Arabidopsis thaliana* (*At*) heterograft which is an exceptional combination of maintainable interfamily grafting (Notaguchi *et al*., 2012; Notaguchi *et al*., 2015). By taking advantage of this finding, we analyzed *Nb* homograft as a compatible model as well as *Nb*/*At* heterograft as an incompatible and more stressful model. We firstly confirmed the more stressful situation in *Nb*/*At* heterograft compared to *Nb* homograft in gene expression level and actual substance transport from the rootstock to the scion. Then, we conducted electron microscopic analysis on the heterografts and found autophagic structures in the cells near the graft boundary. We also conducted microscopy analysis on *At* micrografting and investigated whether the activation of autophagy affects graft wound healing event in *At* and *Nb*.

## Materials and Methods

### Plant materials

The *At* ecotype Columbia (Col) was used as the wild type. *At* and *Nb* seeds were surface sterilized with 5% (w/v) bleach for 5 min, washed three times with sterile water, incubated at 4°C for 3 days, and planted on a medium containing half-strength Murashige and Skoog (1/2 MS) medium, 0.5% (m/v) sucrose, and 1% agar; the pH was adjusted to pH 5.8 with 1 M KOH. Seedlings were grown at 22°C for *At* and 27 °C for *Nb* and 100 µmol m^-2^ s^-1^ of illumination under continuous light conditions.

### Grafting experiments

*Nb*/*At* stem grafting was performed as shown in Okayasu and Notaguchi (2019). Shortly, the stem of 4-week-old *Nb* was wedge grafted on the bolting stem of 5-week-old *At* plant. *Nb* homograft was performed in the same way for *Nb*/*At* heterograft with same aged plants. *At* micrografting was performed using a supportive micro-device, named micrografting chip, developed in Tsutsui *et al*. (2019). Seeds were sown in a prescribed seed pocket of the chip and the chips were put on the Hybond-N+ nylon membrane (GE Healthcare, Chicago, USA) which was placed on the 1/2 MS medium containing 0.5% sucrose and 1% agar. Four-day-old seedlings were conducted to micrografting. In this system, both hypocotyls of stock and scion were cut horizontally with a knife (Kai Co., #2-5726-22 No.11, Tokyo, Japan) and assembled on the chip using forceps. A groove structured on the chip allowed to cut hypocotyl smoothly with the knife and make a fine, flat graft surfaces. Two files of micro-pillars arrayed on the chip supported to hold the hypocotyls of both stock and scion, resulting in uniformly constructed micrografted plants. After grafting, the plants were transferred to a new 1/2 MS medium containing 0.5% sucrose and 2% agar and grown at 27°C for 6 days and then transferred to a new 1/2 MS medium containing 0.5% sucrose and 1% agar and grown at 22°C for 4 days corresponding to 10 days after grafting when the phenotype was examined. The detailed procedures were described in Tsutsui *et al*. (2019). Other *At* sample fractions for hypocotyl partial cutting were prepared using 4-day-old seedlings prepared in the same manner with the plants for micrografting. The *At atg2* mutant line was obtained from the Arabidopsis Biological Resource Center (http://www.arabidopsis.org/abrc). The GFP-ATG8 line used in this study was described in Yoshimoto *et al*. (2004).

### Radio isotope transport assay

Intact *Nb* plants, *Nb* homografts and *Nb*/*At* heterografts were conducted to the radio isotope experiments. All leaves except for three to four pieces of leaves in intact or scion plants were removed, and then the stem was cut at its base and placed in a 5 ml tube that contained 2 ml of distilled water, Pi (0.1 µM), and ^32^P-phosphate (10 kBq). After 6 h incubation at 27°C, the distribution of ^32^P in the plant was visualized by radioluminography using an imaging plate (GE Healthcare UK, Buckinghamshire, UK) and a FLA-5000 image reader (Fujifilm, Tokyo, Japan). The amount of ^32^P was calculated with the image analysis software (Image Gauge version 4.0, Fujifilm). For the time-course analysis, we employed the RRIS (Sugita *et al*., 2016), which enable us to observe ^32^P distribution in the plant sequentially. The activity of ^32^P-phosphate in the incubation solutions was 30 kBq for *Nb* and *Nb*/*Nb* and 60 kBq for *Nb*/*At*. The radioactivity image was captured during the dark period of a 15 min light/dark cycle. The accumulation of ^32^P in the scion and stock was examined in two sections (10×10 cm in size) collected above and below the graft union, respectively.

### VIGS experiments

For virus induced gene silencing (VIGS) of Niben101Scf01320g03007, we called *NbATG5* in this study, a 288-bp portion of the *NbATG5*-coding region was amplified by PCR and the amplified fragment was cloned between *Stu* I and *Mlu* I sites of the CMV-A1 vector (Otagaki *et al.*, 2011). Plasmids containing full-length cDNA of viral RNA were transcribed *in vitro*, and leaves of 3-week-old *Nb* plants were dusted with carborundum and rub-inoculated with the transcripts as described previously (Otagaki *et al.*, 2006). Successful infection of the virus in the upper leaves of *Nb* plants without deletion of inserted sequences was confirmed by RT-PCR of the viral RNA. The sequences of the primers for PCR amplification were shown in Supporting Information Table S1. Primary inoculated leaves were used for secondary inoculation to *Nb* plants. Primary inoculated leaves were ground in 100 mM phosphate buffer (pH 7.0) and 10 µL of the adequate was dropped on the three expanded leaves of a new 3-week-old *Nb* and rub-inoculated. One week after inoculation corresponding to 4-week-old, the infected *Nb* stem was grafted onto the bolting stem of 5-week-old *At* plants. Two weeks after grafting, the successful rate of grafting was scored based on the scion survival.

### Quantitative reverse-transcription polymerase chain reaction

Total RNA was extracted from plant samples using RNeasy MinElute Clean up Kit (Qiagen, Hilden, Germany) following the manufacturer’s instructions. The RNA concentration and quality were determined using NanoDrop One^C^ Spectrophotometer (Thermo Fisher Scientific, Waltham, USA). cDNA synthesis from the RNA sample was carried out using the SuperScript III First-Strand Synthesis SuperMix (Thermo Fisher Scientific) following the manufacturer’s instructions. An equal concentration of cDNA from the samples comparing was used for qRT-PCR reaction with KAPA SYBR Fast qPCR Kit (Sigma-Aldrich, St. Louis, USA). For expression analysis of the genes responsive to nutrient starvation, total RNA was extracted from the graft union of *Nb* scion and *At* stock. For VIGS experiments, total RNA was isolated from the stem of *Nb* of grafted plants 3 days after grafting and used to produce cDNA for qRT-PCR amplification. PCR conditions were 50°C for 2 min, 95°C for 10 min, and 40 cycles of 95°C for 15 s followed by 60°C for 1 min. *NbACT1*, Niben101Scf09133g02006.1, was used as the internal standard. All experiments were performed with three independent biological replicates and three technical replicates. The sequences of the primers were shown in Supporting Information Table S1.

### Microscopy analysis

For observation of resin sections, the graft regions were trimmed by laser blade and fixed with 2% paraformaldehyde and a 2% glutaraldehyde in 0.05 M cacodylate buffer (pH7.4) at 4°C overnight. After this fixation, the samples were washed three times with 0.05 M cacodylate buffer for 30 min each, and were postfixed with 2% osmium tetroxide in a 0.05 M cacodylate buffer at 4°C for 3 h. The samples were dehydrated in graded ethanol solutions (50%, 70%, 90%, and 100%). The schedule was as follows: 50% and 70% for 30 min each at 4°C, 90% for 30 min at room temperature, and four changes of 100% for 30 min each at room temperature. After this dehydration process, the samples were continued to be dehydrated in 100% ethanol at room temperature overnight. The samples were infiltrated with propylene oxide (PO) two times for 30 min each and were put into a 70:30 mixture of PO and resin (Quetol-651, Nisshin EM Co., Tokyo, Japan) for 1 h. Then, the caps of the tubes were opened and PO was volatilized overnight. The samples were transferred to fresh 100% resin and were polymerized at 60°C for 48 h. For light microscopy, the polymerized resins were sectioned at 8 µm with a microtome and mounted on slide glasses. For light microscope observations, sections were stained with 0.5% toluidine blue (pH7.0), mounted on the slide glasses with Mount-Quick (Daido Sangyo Co., Tokyo, Japan), and observed using a digital microscope (DMBA310, Shimadzu RIKA Co., Tokyo, Japan). For transmission electron microscope analysis, the polymerized resins were ultra-thin sectioned at 80–120 nm with a diamond knife using an ultramicrotome (ULTRACUT UCT, Leica, Tokyo, Japan), and the sections were mounted on copper grids. They were stained with 2% uranyl acetate at room temperature for 15 min, and then they were washed with distilled water, followed by a secondary-staining with lead stain solution (Sigma-Aldrich) at room temperature for 3 min. The grids were observed using a transmission electron microscope (JEM-1400Plus, JEOL Ltd., Tokyo, Japan) at an acceleration voltage of 80 kV. Digital images were taken with a CCD camera (VELETA, Olympus Soft Imaging Solutions GmbH, Münster, Germany).

For the analysis of GFP-ATG8 localization, seedlings were treated by ClearSee™ (Fujifilm, Tokyo, Japan) and mounted on glass slides. The slides were examined under a confocal laser scanning microscope system (Zeiss LSM5 PASCAL, Zeiss, Jena, Germany). A 488-nm excitation laser and collecting emission spectrum of 505–530 nm was used for GFP fluorescence detection. For all sample fractions, more than 3 independent samples were observed.

## Results

### Grafting caused nutrient deficient situation

To investigate events during grafting, we firstly analyzed two graft combinations, *Nb*/*Nb* compatible homograft and *Nb*/*At* less compatible heterograft. We investigated the level of stress caused by graft wound in these grafts by examining the extent of mass flow from the stock to the scion and gene expression at graft junctions (Fig. 1). To examine the mass transport from the stock to the scion, we applied a radio isotope species, ^32^P, to the stem of the stock plants 14 days after grafting. The stem of the stock was cut off below the grafting point and soaked into a solution containing Pi labeled with ^32^P for 2 hours to incorporate Pi. Intact *Nb* plant was also treated in the same way as a control experiment. The stems incorporating the radioactive label were directly brought into contact with the imaging plate to expose the image, and the dose were measured with a scintillation image detector over time (Supporting information Fig. S1). Although the isotope signal was detected from the intact *Nb* stem and the scion parts of both grafts, the signal in the scion part was faint and intense signal of isotope was detected just below the grafting point in the *Nb*/*At* heterograft, suggesting that the upward mass transport was restricted in the *Nb*/*At* heterograft (Fig. 1a). Fig. 1(b) shows comparisons of the measured values divided by the projected area on the scion and stock part of the graft. For intact plants, we split the stems where appropriate and took the ratios of the top and bottom as well. As a result, it was found that the ratio in the *Nb*/*At* heterograft was smaller compared to that in intact *Nb* plants. In the intact *Nb* plant, the ratio of the value of scion part including stem, leaves and apical meristem to that of stock stem was approximately 1.8, whereas in the *Nb*/*At* heterograft the ratio was about 0.3. These differences were significant at the 1% level by Student’s *t* test. In contrast, the ratio in the *Nb*/*Nb* homograft was approximately 1.9, which was not clearly different from that in the intact plants. These results suggested that the transport of Pi was fully restored in homograft 14 days after grafting. On the other hand, the transport of Pi was small in heterogeneously grafted plants, suggesting much stressful conditions were continued even at this point in time (Fig. 1b).

**Fig. 1.**
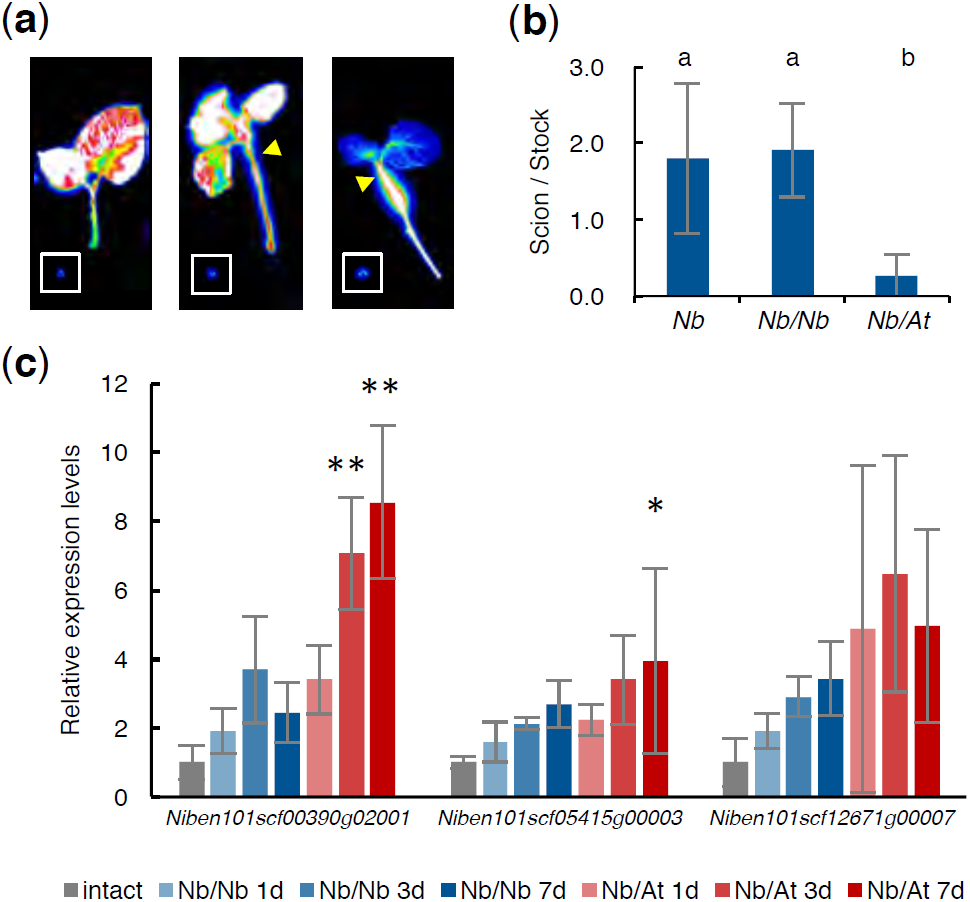
Grafting caused nutrient deficient situation. (a) Phosphorus transport in an intact *Nb* plant, a *Nb*/*Nb* homograft and a *Nb*/*At* heterograft from left to right. All plantlets were incorporated inorganic phosphate (Pi) labeled with ^32^P from the cut stems. The signal amount is shown in heat map; from white to blue corresponding to from higher to lower. Insets indicate the reference points to compare the intensity. Arrowheads indicate the grafted points. (b) The quantification of the transport of Pi by imaging plates at 6 h after application. Differences between the sample groups were tested by a two-way ANOVA followed by a Tukey’s post hoc tests, with α set at *P* < 0.01 (biological replicates; n=11– 13). (c) Relative expression of the genes responds to nutrient starvation. The expression levels of *Nb* genes were established by real-time PCR analysis in the intact *Nb, Nb*/*At* and *Nb*/*Nb* grafted samples. Expression levels were normalized by the levels of *NbACT1* and adjusted to be relative to the intact samples. RNA was extracted from the stem tissues of 10 intact or grafted plants for each sample fractions. Mean ± s.d. from 3 independent experiments. Statical analyses were performed by ANOVA with Dunnett’s multiple comparison of means test; *P* < 0.05 (*) or *P* < 0.01 (**).

We considered that the grafted scions were subject to inorganic nutrient starvation stress before forming the vasculature connection between the scion and stock, and examined the expression level of marker genes that respond to mineral nutrient starvation stress. We focused on the *ERD5, SGT1*, and *NRT2.5* genes of *At* among the nutrient deficiency responsive genes, reported previously (Hammond *et al*., 2003; Kiba *et al*., 2018), and identified their orthologs in *Nb* by tblastx search. The orthologs of *Nb* of *AT3G30775* (*ERD5*), *AT2G43820* (*SGT1*) and *AT1G12940* (*NRT2.5*) were *Niben101Scf00390g02001, Niben101Scf05415g00003* and *Niben101Scf12671g00007*, respectively. The expression of these three genes on the 1, 3 and 7 days after grafting and the expression in intact plants were quantified by quantitative reverse transcription PCR (Fig. 1c). The *Nb*/*Nb* homograft sample showed an increasing tendency of the gene expression 1 and 3 days after grafting. The degree of increase for all three genes was larger in the *Nb*/*At* heterografts. It gradually increased from 1 to 7 days after grafting, again showing that the heterografts have much stressful condition compared to the homografts. A tendency of this nutrient deficiency response was also observed in the *At* stock part of the *Nb*/*At* heterografts, however significant difference was not detected in our statical analysis (Supporting information Fig. S2).

### Microautophagy-like reactions occurred at the graft boundary

To see the cellular event during graft healing processes, we analyzed graft junction of the *Nb*/At heterograft which would exert reaction to withstand stressful situation after grafting more significantly than less stressful compatible grafts. The cross sections of the grafted junction of the *Nb*/At heterograft 2 weeks after grafting were observed with an optical microscope and a transmission electron microscope (TEM) (Fig. 2). At this time point, the graft surfaces were sufficiently bounded, but the boundary surface could be identified relatively clearly. In particular, callus-like cells were formed at the boundary around the cambial region (Fig. 2b–e). We noticed that there were many granular structures in both *Nb* and *At* cells near the graft boundary surfaces where *Nb* cells and *At* cells are in contact together. Such structure was not found in normal cells. These granular structures were observed as if they penetrated from the vacuole membrane or floated in the vacuole (Fig. 2c,e,f). In high magnification images, various types of granules were found, including granules of texture similar to the cytoplasmic matrix, mitochondrial cristae-like structures, and plastid containing grana structures (Fig. 2f–i). These seem to be typical microautophagy like structures. Based on previous reports that nutrient starvation induces autophagy (Bassham *et al*., 2006), we hypothesized that autophagy is induced by the wound stress including nutrient deficient stress at the grafting region.

**Fig. 2.**
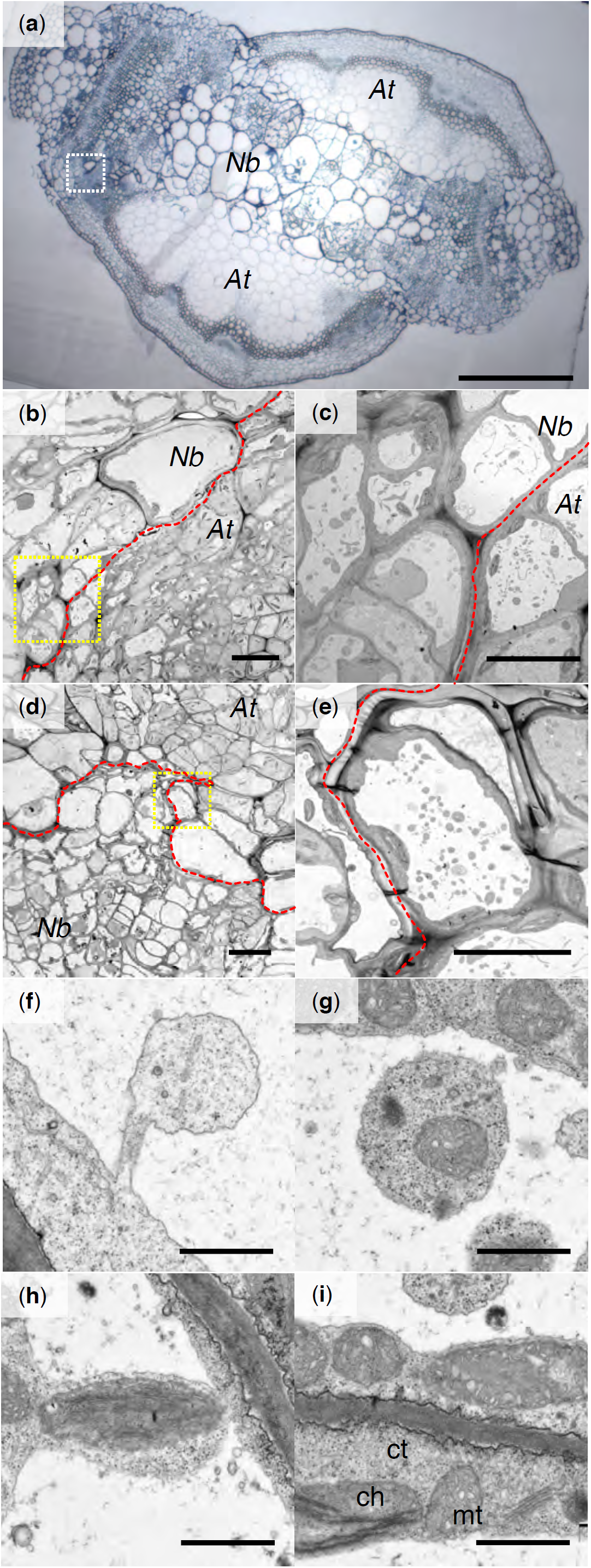
Observation of the cells near *Nb*/*At* heterograft boundary 14 days after grafting. (a) Optical microscopic image of a horizontal slice of the grafted region. Dashed rectangle indicates the area of (b). Bar = 500 µm. (b)–(i) TEM images of the *Nb*/*At* heterografts. (b) Graft boundary around the region where the both cambial tissues of the scion and the stock plants located close. A dashed rectangle coloured in yellow indicates the area of (c). Bar = 20 µm. (c) A magnified image of the boundary cells. Bar = 10 µm. (d) Another boundary of the *Nb*/*At* heterograft. A sashed rectangle coloured in yellow indicates the area of (e). Bar = 20 µm. (e) A magnified image of the *Nb* boundary cells in (d). Slashed lines coloured in red indicate graft boundaries. Bar = 10 µm. (f)–(h) Magnified images of characteristic granular structures in the vacuoles of the boundary cells. Bars = 1 µm. (i) A magnified image of the cytosolic region of the boundary cell. Bar = 1 µm. ‘mt’, ‘ch’ and ‘ct’ indicate mitochondria, chloroplast and cytosol, respectively.

So, in this study, we observed the localization of autophagy-related components using a GFP-fusion marker line in order to verify whether autophagy is induced at the grafted sites. ATG8 is normally expressed ubiquitously in cells, and it is known that when autophagy mechanism is activated, they are localized in autophagosome and endosomal microautophagy machinery (Izumi *et al*., 2017). We performed hypocotyl micrografting using an *At* GFP-ATG8 transgenic line which constitutively expresses the GFP fused-ATG8 protein under the CaMV 35S promoter, and observed the grafted junction region with a confocal laser microscope. In the homografts of the GFP-ATG8 transgenic line, the fluorescence of GFP-ATG8 was observed 3 day after grafting. The fluorescence signals were especially detected in the scion area above the graft surface, and weaker in the stock (Fig. 3a,e,i). To clarify the location of autophagy activation, we performed reciprocal grafting between the GFP-ATG8 line and wild type (WT). As results, the fluorescence of GFP-ATG8 was only detected clearly when the GFP-ATG8 line was grafted as the scion side, but the signals were very faint in the reverse grafting (Fig. 3b,c,f,g,j,k). These results suggest that the autophagy machinery was induced in the grafted region, especially above the grafted surface where the stress level may be high (Figs. 1c, S2). More detailed observation of the site where the GFP fluorescence was detected often resulted in the signal appearing as dots (Fig. 3a,b,e,f). Interesting notion here is that, when the GFP-ATG8 line was grafted as a scion to the WT rootstock, a fluorescence signal was also observed in the WT rootstock below the graft surface (Fig. 3b, f, j), which was distinguishable from the autofluorescence detected in the vascular structures and in the vicinity of the cut edge in the WT (Fig. 3d,h). This GFP signal may indicate that GFP-ATG8 proteins moved from the scion to the stock to some extent, like as the translocation of organelle maker proteins over graft junction previously observed (Paultre *et al.*, 2016). The fluorescence of these GFP-ATG8 was rarely observed 7 days after grafting (Fig. 3l), except when the grafts were not sufficiently fused (data not shown).

**Fig. 3.**
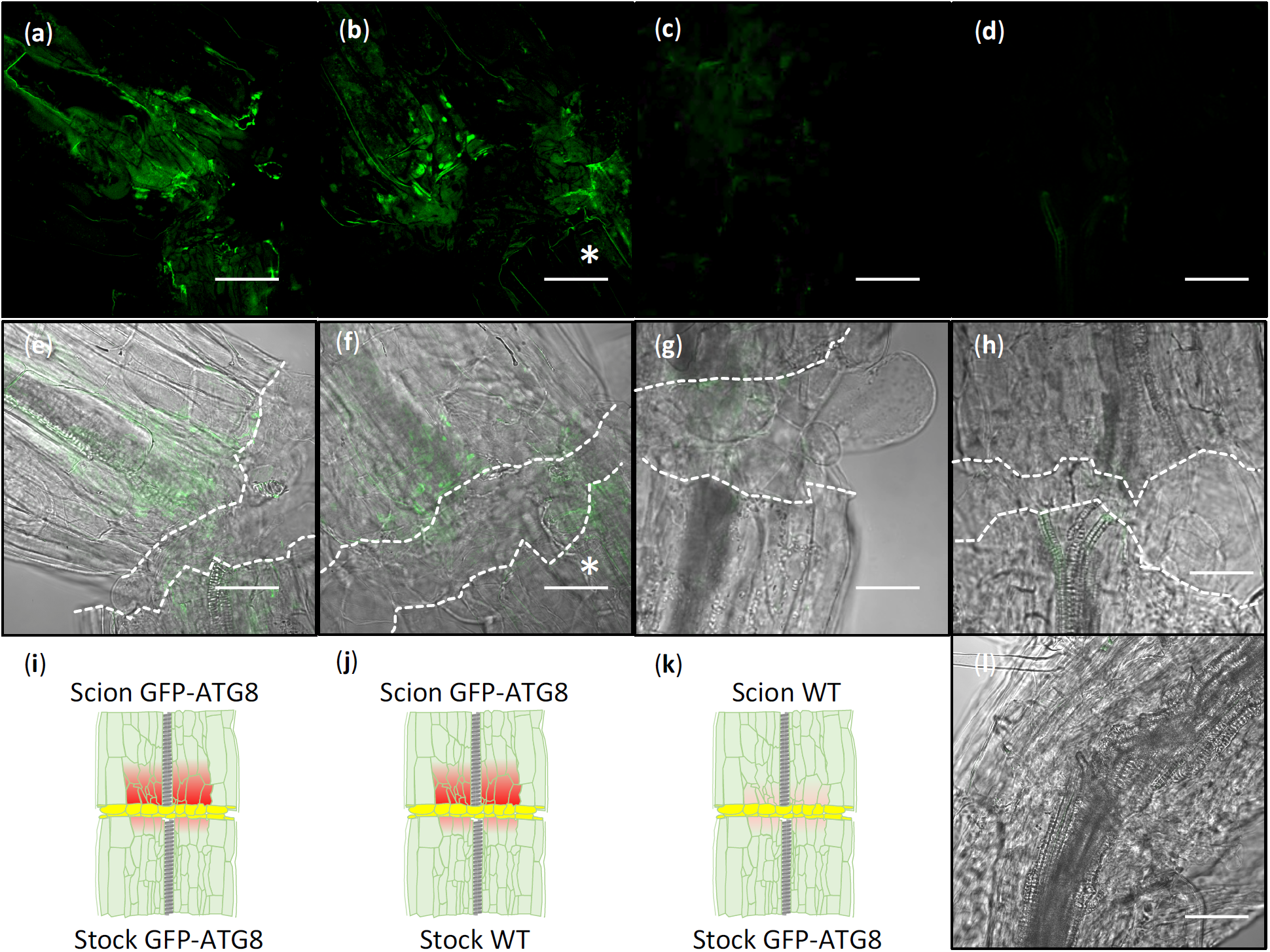
Autophagy was induced at the graft boundary of the scion side in the *At* homograft. (a)–(d) Representative GFP fluorescent images detected by confocal laser scanning microscope in the graft junctions of the *At* wild type and transformant constitutively expressing GFP-ATG8 at 3 days after grafting. (e)–(h) Blight frame (DIC) and the fluorescence frame merged images. Slashed lines indicate the cutting edges of grafting. Asterisks in (b), (f) indicate the fluorescence signal detected in the stock part. (i)–(k) Illustrated images of the graft samples and the GFP-ATG8 detected area (coloured in red). Yellow area indicates callus tissues formed between the scion and stock plants. (a), (e), (i) GFP-ATG8/GFP-ATG8, (b), (f), (j) GFP-ATG8/WT, (c), (g), (k) WT/GFP-ATG8, (d), (h) WT/WT grafts. (l) Blight frame and the fluorescence frame merged image of the GFP-ATG8/GFP-ATG8 graft 7 days after grafting. Bars = 50 µm.

### Autophagy was induced by the cutting

It is known that ATG8 protein is constitutively expressed but does not exhibit its function under normal conditions and only when the autophagy is activated, ATG8 protein aggregates within the autophagy mechanism. We confirmed that constitutively expressed GFP-ATG8 was dispersed in the hypocotyl of the intact transgenic plants and was not detected as a fluorescent signal (Fig. 4a,d,g). Then, we verified the possibility that autophagy was activated in the case of wounding repair (Fig. 4c–f,h,i). The hypocotyls of the transgenic plants were cut in two ways, a half-cut and a full-cut. As a result, upon repairing the injury, the fluorescence signals of GFP-ATG8 were detected in the region around the half-cut wounded site, particularly in the region excluding the epidermis and endothelium above the wounded site. In contrast, the GFP-ATG8 fluorescence signal was not clearly detected at the cutting end of the full-cut hypocotyls which did not contact to the other side of wounds. Callus induction was observed at the cut surface of the full-cut hypocotyls similar to the cases of grafting and half-cut wound repair. This provoked an insight that autophagy is only activated when the wounded part faces to the encountering wounded part.

**Fig. 4.**
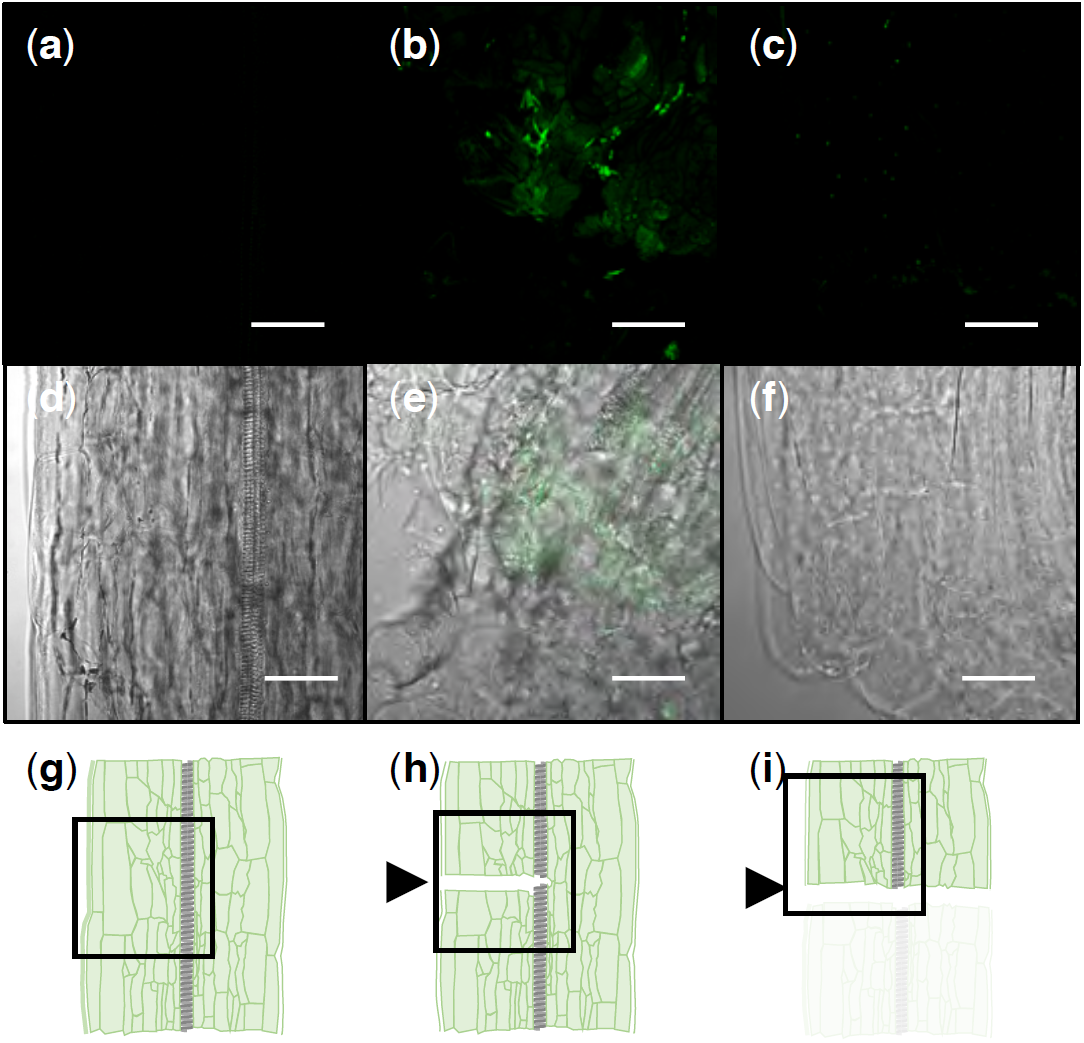
Autophagy was induced during wound repairing, but not induced by cutting without contacting to the other side of wounds. (a)–(c) Representative GFP fluorescent signals detected in the GFP-ATG8 plants by confocal laser scanning microscope. (d)–(f) Blight flame (DIC) and the fluorescence frame merged images. (g)–(i) Illustrated images of the sample processing. Rectangles indicate the area detecting the fluorescence and blight flame images in (a)–(f). Arrowheads indicate cut positions. (a), (d), (g) An intact hypocotyl, (b), (e), (h) a half-cut hypocotyl and (c), (f), (i) upper part of a full-cut hypocotyl of the GFP-ATG8 plants 3 days after incision. Bars = 50 µm.

### Autophagy contributed to wound healing in homograft and cutting

In order to verify whether the mechanism of autophagy is critical for the establishment of grafts and wound repair, graft or wound healing experiments were performed using a mutant of an autophagy-related gene, *ATG2*. The young seedlings of *atg2* mutant showed no phenotype under normal growth conditions, and the growth of *atg2* mutant plants was equivalent to the WT plants when measured the fresh weight of the shoot part (Fig. 5a–c). In the successful rates of the grafting, there was no significant difference among *atg2* mutant and WT self-grafts in four independent grafting experiments (Supporting Information Table S2). However, comparing the fresh weight of successfully grafted plants 10 days after grafting, it was seen a difference. The fresh weights of *atg2* mutant self-grafts were significantly less than that of WT self-grafts in every experiment, suggesting *atg2* mutation affected the growth of the scion shoots (Fig. 5d–f). It provided an evidence that autophagy has a partial role on the efficiency of the graft wound healing. In addition, when the hypocotyl was partly cut, around a half of the hypocotyl to mimic the wound situation, the fresh weights of the shoot part of *atg2* mutant plants were significantly less than those of WT plants (Fig. 5g–i). Taken together, these results indicate that the autophagy mechanism is important for healing of naturally occurring wounds as well as the artificial graft wounds.

**Fig. 5.**
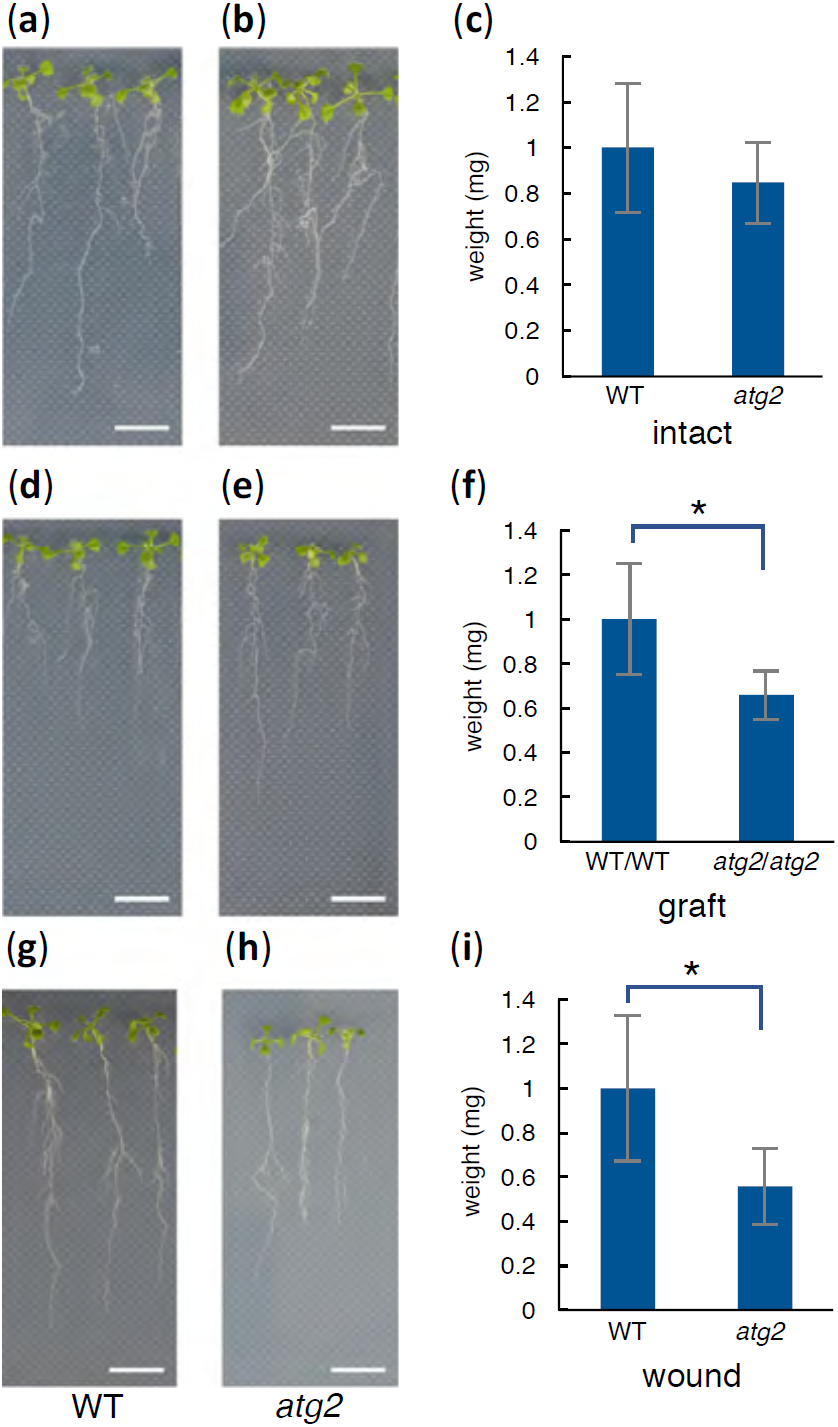
Autophagy contributed to wound healing in homograft and cutting in *At*. Images of 7-day-old intact wild type plants (a) and *atg2* mutant intact plants (b). (c) Comparison of the shoot fresh weight of the intact wild type and *atg2* mutant plants 7 days after germination (12 seedlings were conducted for each sample fraction). Images of wild type self-grafts (d) and *atg2* self-grafts (e) 10 days after grafting. (f) Comparison of the shoot fresh weight of the wild type and *atg2* mutant self-grafts 10 days after grafting (see also Supporting Information Table S2). Images of wounded wild type (g) and *atg2* mutant plants (h) 10 days after cutting. (i) Comparison of the shoot fresh weight of the wounded wild type and *atg2* mutant plants 10 days after cutting. Experiments were performed three times. For each experiment, 15–18 seedlings were conducted for each sample fraction. Bars = 10 mm. Student’s *t* tests were conducted, *P* < 0.05 (*).

### Autophagy contributed to wound healing in heterograft

So far, we have shown the involvement of autophagy in the adhesion of grafts based on the granular structures in the vacuole of cells at the boundary found in TEM images of *Nb*/*At* heterografts. In order to verify whether such an autophagy mechanism is necessary to establish grafting between *Nb* and *At* plants, we targeted *NbATG5* which is the most similar gene to *AtATG5* (AT5G17290) in *Nb* searched by tblastx. A *Nb* plant which *NbATG5* gene was knocked down by CMV VIGS system (*NbATG5* VIGS) (Otagaki *et al.*, 2011) were grafted to an *At* bolting stem and measured the survival rate 14 days after grafting. As control experiments, no infected (NI) or infected with a CMV expressing a part of GFP sequence *Nb* plants (*GFP* VIGS) were also grafted to *At* stock plants. The procedures of our knock down experiments are shown in Fig. 6(a). First, we confirmed the suppression of *NbATG5* by VIGS. The expression levels of *NbATG5* gene were quantified at the grafted region of the *Nb*/*At* heterografts of the NI, *GFP* VIGS and *NbATG5* VIGS 3 days after grafting. In the heterografts of the *NbATG5* VIGS, the *NbATG5* expression was significantly decreased compared to the heterografts of the NI or *NbATG5* VIGS (Fig. 6b). Then the survival rate of these grafts was examined at 14 days after grafting. The survival rate of the *NbATG5* VIGS heterografts was also significantly reduced compared to the heterografts of the NI or *NbATG5* VIGS (Fig. 6c). These results suggest that *NbATG5* gene plays a role in the establishment of heterologous grafts between *Nb* and *At* plants.

**Fig. 6.**
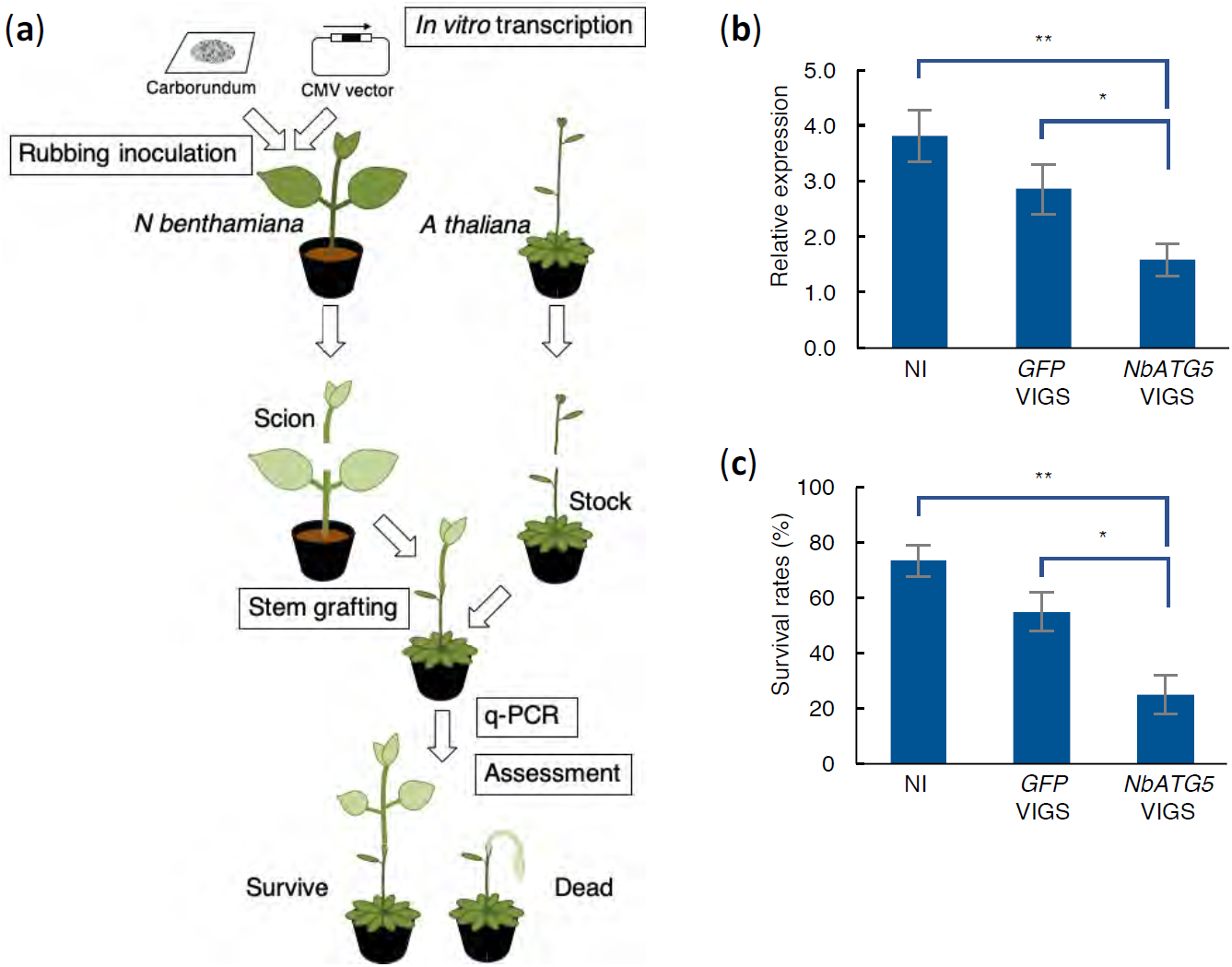
Autophagy contributed to wound healing in heterograft. (a) The scheme for the *Nb*/*At* grafting experiment with the VIGS treatment. (b) Suppression of *NbATG5* gene expression by the VIGS was verified by qRT-PCR (3 grafts were conducted for each sample fraction). Expression levels were normalized by the levels of *NbACT1* and adjusted to be relative to the sample with no virus infection (NI). (c) Effect of suppression of *NbATG5* gene expression by the VIGS on graft establishment. Experiments were performed three times. For each experiment, 10 grafts were conducted for each sample fraction. Differences between the sample groups were tested by a two-way ANOVA followed by a Tukey’s post hoc tests, with α set at *P* < 0.05 (*) or *P* < 0.01 (**).

## Discussions

Graft wounds cause drastic physiological changes to withstand the injury stress. The grafted wound sites are repaired by a sequence of processes such as primary wound response, callus formation, cell adhesion, tissue differentiation and vascular formation. This healing processes occur within a short period, generally several days to a week, in herbaceous compatible grafts making the detailed study of grafting difficult. Graft compatibility is related to genetic distance of the plants and grafting is generally performed in same species/genus/family. However, a few maintainable interfamily grafts have been reported (Nickell, 1948; Kollmann & Glockmann, 1985; Notaguchi *et al*., 2012) and they showed less mass flow among the stock and scion (Rachow-Brandt & Kollmann, 1992; Fig. 1a,b) and higher stressful response in gene expression (Fig. 1c). Thus, graft compatibility affects the efficiency of repairing at the graft wounds. This study took advantage of less compatible grafts to explore the undetermined cellular events how to overcome the stressful wound conditions at grafted sites and identified activation of autophagy, as a cellular event occurred at the graft wounds. Activation of autophagy was observed in *Nb*/*At* heterografts and *At*/*At* homografts (Figs. 2, 3). Not only grafting situation, the autophagy was also activated when plants got injury on the stem (Fig. 4). The knock-down or knock-out of autophagy components retarded the recovery from graft or wound incision, suggesting a positive role of autophagy in these wound repairing (Figs. 5, 6). Still the clear link among autophagy activation and the other cellular processes during graft establishment are missing, but autophagy is activated to function in graft healing process when the cut surface is in contact with another cut surface (Fig. 7).

**Fig. 7.**
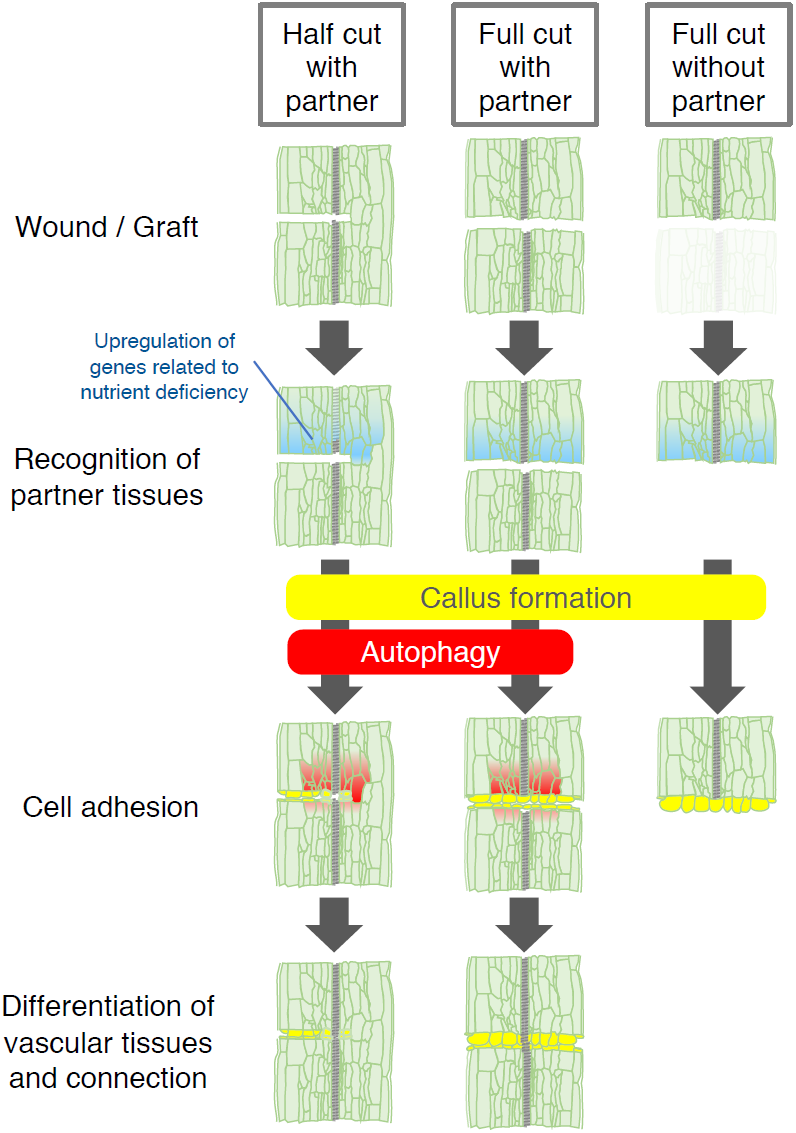
A model for wound healing on grafting and cutting. In grafted or wounded plants, the vascular rupture near the wound sites results in a starvation of mineral nutrition. Mineral nutrition starvation is more severe at the top of the wound sites than at the bottom. Callus formation is induced from the cut surface. Autophagy is activated when the wound site is contact with the other wound site, but not without wounds encountering. Finally, wounds or grafted sites are healed through cell adhesion and differentiation of vascular tissues and connection.

Autophagy is known as a nutrient recycling reaction triggered by nutrition starvation in plants. In this study, we observed a number of granular structures in the vacuoles of cells near the graft junction, that were not found in other cells, when we observed the graft plant tissues of *Nb*/*At* heterograft samples using TEM (Fig. 2c,e–h). Because some structures were found in which the vacuolar membrane took in the cytoplasm by engulfing, it was speculated that microautophagy was occurring. It is known that there are at least three types of autophagy. In macroautophagy, it is thought that after the double layered autophagosomes is formed, the vacuole membrane and the outer membrane of autophagosome fuse to incorporated the granules into the vacuole. When chloroplasts are taken up by autophagosomes, it is thought that the grana structure is not completely taken in, but a part of the stroma is cut off and taken up, and gradually undergoes digestion by vacuole (Ishida *et al*., 2008). In contrast, in the process of microautophagy named chlorophagy, it has been reported that the entire chloroplast, including the grana structure, is collectively incorporated into the vacuolar membrane (Izumi *et al*., 2019; Nakamura & Izumi, 2019). We observed that the structure of chloroplast was taken into the vacuole with the whole membrane structure, therefore, the phenomenon we observed in this case was appeared to be a type of microautophagy. Previous study on a heterograft between *Sedum telephoides* and *Solanum penellii* reported done by Moore & Walker (1981) reported the occurrence of cytoplasmic vesiculation and the loss of cellular membrane integrity in the cells of *Sedum*. Such cellular necrosis was sometimes observed in our experiments but was different from the autophagy granule locating in vacuoles.

Although the degree of induction of nutrient deficiency response genes was smaller in homografts than heterografts in the case of *Nb* grafts, we determined that autophagy was similarly induced in homografts using the *At* GFP-ATG8 autophagy maker line. Moreover, in the grafted samples 3 days after grafting, remarkable GFP-ATG8 expression was detected on the scion side above the grafted site (Fig. 3a,b,e,f). This correlates with the fact that the expression of the nutrient deficiency responsive gene was significantly increased on the scion side (Figs. 1c, S2), suggesting that the autophagy is mainly functions in the scion wound sites.

Grafting is an artificial event, and in particular heterografting is unlikely to occur in nature. Therefore, we thought that the original physiological significance of autophagy observed during grafting would appeared in wound repairing. After the cut or graft wound was treated on the hypocotyl of the *At* plants, the fluorescence of GFP-ATG8 was detected around the wound sites 3 days after wound treatment. Interestingly, no signal was detected when the wounded surface was placed without contact to another wound site. These results may mean that the induction of autophagy at the wounds occurs only when the tissue recognizes the presence of contactable or proximate, healable confronting tissue. Furthermore, since callus induction occurs even in the wound site without confronting tissue, induction of autophagy during wound healing may be an independent event from callus formation (Fig. 7).

In this study, we examined the effect of mutant or knockdown of the genes encoding essential autophagy components on graft establishment using *At* self-graft and *Nb*/*At* heterograft. As a result, the success rate of self-graft did not decrease, whereas the success rate of heterograft significantly decreased. In the *At* self-grafts, the occurrence of autophagy was detected only for a short period immediately after grafting; it was detected 3 days after grafting, but was hardly detected 7 days after grafting. However, even in the self-grafts, there was a significant difference in the fresh weight of the scion plants after grafting. Although autophagy is not essential for the establishment of grafts, it is possible that if there is affinity or compatibility for grafts, it acts on the nursing of tissues after graft incision, helping to recover from the injury. On the other hand, a large number of microautophagy granules were observed on the grafted surface of the heterograft 14 days after grafting, suggesting that the autophagy phenomenon has continued for a long time in the less compatible heterografts. These results suggest that microautophagy functions under conditions of lower affinity or severe conditions for the establishment of grafts, and supports the maintenance of the grafts.

Grafting achieved through many of key cellular/tissue level events, including rapid wound response, nutrient and hormonal regulation, cell regeneration and tissue reunion through cell differentiation for each requiring cell types. Hence, a future question would be which process(es) is regulated by autophagy machinery and which is not. As the wound repairing is essential for plants to survive robustly in nature, several independent pathways could be functioning during wound healing process. By taking advantage of availability of autophagy mutants, potential autophagy related processes accelerated by wounding/grafting can be addressed in future studies. Studying on wounding/grafting will gain insights into the biological function of autophagy in plants.

## Acknowledgements

We thank M. Hattori, M. Matsumoto, I. Yoshikawa and A. Yagi for technical assistance. We are grateful to K. Yoshimoto (Meiji University, Japan) for providing *At* seeds of GFP-ATG8 and *atg2*, C. Masuta (Hokkaido University, Japan) for providing CMV vectors and S. Otagaki (Nagoya University, Japan) for advising VIGS experiments. This work was supported by grants from Japan Society for the Promotion of Science Grants-in-Aid for Scientific Research (18KT0040 and 19H05361), and the Cannon Foundation (R17-0070) to M.N.

## Conflict of interest

We declare no conflict of interest.

## Author contributions

KK and MN conceived of the research and designed experiments. YK, RT and KO performed grafting experiments, RS and KT performed radio isotope experiments, RT performed VIGS experiments and KK performed microscopy analysis. KK and MN wrote the paper.

## Supporting Information

**Fig. S1** Radio isotope transport experiments in intact *Nb, Nb*/*Nb* homograft and *Nb*/*At* heterograft.

**Fig. S2** Relative expression of the genes related to nutrient deficiency in the *At* stock part of *Nb*/*At* heterografts.

**Table S1** The sequences of the PCR primers used in this study.

**Table S2** Comparisons of the success rates of *At* wild type and *atg2* mutant self-grafts.

## References

Asahina M, Azuma K, Pitaksaringkarn W, Yamazaki T, Mitsuda N, Ohme-Takagi M, Yamaguchi S, Kamiya Y, Okada K, Nishimura T, Koshiba T, Yokota T, Kamada H, Satoh S. 2011. Spatially selective hormonal control of RAP2.6L and ANAC071 transcription factors involved in tissue reunion in Arabidopsis. Proceedings of National Academy of Science of the United States of America 108: 16128–16132.

Asahina M, Iwai H, Kikuchi A, Yamaguchi S, Kamiya Y, Kamada H, Satoh S. 2002. Gibberellin produced in the cotyledon is required for cell division during tissue reunion in the cortex of cut cucumber and tomato hypocotyls. Plant Physiology 129:201–210

Asahina M, Satoh S. 2015. Molecular and physiological mechanisms regulating tissue reunion in incised plant tissues. Journal of Plant Research 128: 381–388.

Assunção1 M, Santos C, Brazão J, Eiras-Dias JE, Fevereiro P. 2019. Understanding the molecular mechanisms underlying graft success in grapevine. BMC Plant Biology 19: 396.

Bassham DC, Laporte M, Marty F, Moriyasu Y, Ohsumi Y, Olsen LJ, Yoshimoto K. 2006. Autophagy in development and stress responses of plants. Autophagy 2: 2–11.

Chanoca A, Kovinich N, Burkel B, Stecha S, Bohorquez-Restrepo A, Ueda T, Eliceiri KW, Grotewold E, Otegui MS. 2015. Anthocyanin vacuolar inclusions form by a microautophagy mechanism. Plant Cell 27: 2545–2559.

Chen Z, Zhao J, Hu F, Qin Y, Wang X, Hu G. 2017. Transcriptome changes between compatible and incompatible graft combination of Litchi chinensis by digital gene expression profile. Scientific Reports 7: 3954.

Davies Jr. FT, Geneve RL, Wilson SB. 2018. Hartmann & Kester’s Plant Propagation: Principles and Practices, 9th edn. Pearson, NY, 490–542.

van Doorn WG, Papini A. 2013. Ultrastructure of autophagy in plant cells: a review. Autophagy 9: 1922e1936.

Farré JC, Subramani S. 2016. Mechanistic insights into selective autophagy pathways: lessons from yeast. Nature Reviews Molecular Cell Biology. 17: 537–552.

Goldschmidt EE. 2014. Plant grafting: new mechanisms, evolutionary implications. Frontiers in Plant Science 5: 727.

Gómez-Sánchez R, Rose J, Guimarães R, Mari M, Papinski D, Rieter E, Geerts WJ, Hardenberg R, Kraft C, Ungermann C et al. 2018. *Atg9* establishes *Atg2*-dependent contact sites between the endoplasmic reticulum and phagophores. Journal of Cell Biology 217: 2743–2763.

Hammond J, Bennett MJ, Bowen HC, Broadley MR, Eastwood DC, May ST, Rahn C, Swarup R, Woolaway KE, White PJ. 2003. Changes in Gene Expression in Arabidopsis Shoots during Phosphate Starvation and the Potential for Developing Smart Plants. Plant Physiology 132: 578–596.

Ishida H, Yoshimoto K, Izumi M, Reisen D, Yano Y, Makino A, Ohsumi Y, Hanson MR, Mae T. 2008. Mobilization of Rubisco and Stroma-Localized Fluorescent Proteins of Chloroplasts to the Vacuole by an ATG Gene-Dependent Autophagic Process. Plant Physiology 148: 142–155.

Izumi M, Ishida H, Nakamura S, Hidema J. 2017. Entire photodamaged chloroplasts are transported to the central vacuole by autophagy. Plant Cell 29: 377–394.

Izumi M, Nakamura S, Li N. 2019. Autophagic Turnover of Chloroplasts: Its Roles and Regulatory Mechanisms in Response to Sugar Starvation. Frontiers in Plant Science. 10: 280.

Jeffree CE, Yeoman MM. 1983. Development of intercellular connections between opposing cells in a graft union. New Phytology 93: 491–509.

Kiba T, Inaba J, Kudo T, Ueda N, Konishi M, Mitsuda N, Takiguchi Y, Kondou Y, Yoshizui T, Ohme-Takagi M et al. 2018. Repression of Nitrogen-Starvation Responses by Members of the Arabidopsis GARP-Type Transcription Factor NIGT1/HRS1 Subfamily. Plant Cell 30: 925–945.

Kollmann R, Glockmann C. 1985. Studies on graft unions. I. Plasmodesmata between cells of plants belonging to different unrelated taxa. Protoplasma 124: 224–235.

Kollmann R, Yang S, Glockmann C. 1985. Studies on graft unions II. Continuous and half plasmodesmata in different regions of the graft interface. Protoplasma 126: 19–29.

Krick R, Muehe Y, Prick T, Bremer S, Schlotterhose P, Eskelinen EL, Millen J, Goldfarb DS, Thumm M. 2008. Piecemeal microautophagy of the nucleus requires the core macroautophagy genes. Molecular Biology of Cell 19: 4492–4505.

Kwon SI, Cho HJ, Jung JH, Yoshimoto K, Shirasu K, Park OK. 2010. The Rab GTPase RabG3b functions in autophagy and contributes to tracheary element differentiation in Arabidopsis. The Plant Journal. 64: 151–164.

Li F, Vierstra RD. 2012. Autophagy: a multifaceted intracellular system for bulk and selective recycling. Trends in Plant Science. 17: 526–537.

Liu Y, Bassham DC. 2012. Autophagy: pathways for self-eating in plant cells. Annual Reviews of Plant Biology. 63: 215–237.

Lulu Xie, Chunjuan Dong and Qingmao Shang. 2019. Gene co-expression network analysis reveals pathways associated with graft healing by asymmetric profiling in tomato. BMC Plant Biology 19: 373.

Maathuis FJM. 2009. Physiological functions of mineral macronutrients. Current Opinion on Plant Biology 12: 250–258.

Matsuoka K, Sugawara E, Aoki R, Takuma K, Terao-Morita M, Satoh S, Asahina M. 2016. Differential Cellular Control by Cotyledon-Derived Phytohormones Involved in Graft Reunion of Arabidopsis Hypocotyls. Plant and Cell Physiology 57: 2620–2631

Melnyk CW, Gabel A, Hardcastle TJ, Robinson S, Miyashima S, Grosse I, Meyerowitz EM. 2018. Transcriptome dynamics at Arabidopsis graft junctions reveal an inter tissue recognition mechanism that activates vascular regeneration. Proceedings of National Academy of Science U.S.A. 115: E2447–E2456.

Melnyk CW, Schuster C, Leyser O, Meyerowitz EM. 2015. A Developmental Framework for Graft Formation and Vascular Reconnection in *Arabidopsis thaliana*. Current Biology 25: 1306–1318.

Moore R, Walker DB. 1981. Studies of vegetative compatibility-incompatibility in higher plants. I. A structural study of a compatible autograft in *Sedum telephoides (Crassulaceae*). American Journal of Botany 68: 820–830.

Nakamura S, Hidema J, Sakamoto W, Ishida H, Izumi M. 2018. Selective elimination of membrane-damaged chloroplasts via microautophagy. Plant Physiology. 177: 1007–1026.

Nakamura S, Izumi M. 2019. Chlorophagy is ATG gene-dependent microautophagy process. Plant signaling & behavior 14: 1554469.

Nickell LG. 1948. Heteroplastic Grafts. Science 108: 389.

Otagaki S, Arai M, Takahashi A, Goto K, Hong JS, Masuta C, Kanazawa A. 2006. Rapid induction of transcriptional and post-transcriptional gene silencing using a novel *Cucumber mosaic virus* vector. Plant Biotechnology 23: 259–265.

Otagaki S, Kawai M, Masuta C, Kanazawa A. 2011. Size and positional effects of promoter RNA segments on virus-induced RNA-directed DNA methylation and transcriptional gene silencing. Epigenetics 6: 681–691.

Paultre D, Gustin MP, Molnar A, Oparka KJ. 2016. Lost in transit: long-distance trafficking and phloem unloading of protein signals in Arabidopsis homografts. Plant Cell 28: 2016–2025.

Pitaksaringkarn W, Ishiguro S, Asahina M, Satoh S. 2014. ARF6 and ARF8 contribute to tissue reunion in incised Arabidopsis inflorescence stems. Plant Biotechnology 31: 49–53.

Rachow-Brandt G, Kollmann R. 1992. Studies on graft unions IV. Assimilate transport and sieve element restitution in homo- and heterografts. Journal of Plant Physiology 139: 579–583.

Sala K, Karcz J, Rypień A, Kurczyńska EU. 2019. Unmethyl-esterified homogalacturonan and extensins seal Arabidopsis graft union. BMC Plant Biology 19: 151.

Soto-Burgos J, Zhuang X, Jiang L, Bassham DC. 2018. Dynamics of autophagosome formation. Plant Physiology. 176: 219–229.

Tsutsui H, Yanagisawa N, Kawakatsu Y, Ikematsu S, Sawai Y, Tabata R, Arata H, Higashiyama T, Notaguchi M. 2019. Micrografting device for testing environmental conditions for grafting and systemic signaling in Arabidopsis. bioRxiv doi: 10.1101/2019.12.20.885525.

Wang H, Zhou P, Zhu W, Wang F. 2019. De novo Comparative Transcriptome Analysis of Genes Differentially Expressed in the Scion of Homografted and Heterografted Tomato Seedlings. Science Reports 9: 20240.

Wang YQ. 2011. Plant grafting and its application in biological research. Chinese Science Bulletin 56: 3511–3517.

Yoshimoto K, Hanaoka H, Sato S, Kato T, Tabata S, Noda T, Ohsumi Y. 2004. Processing of ATG8s, Ubiquitin-Like Proteins, and Their Deconjugation by ATG4s Are Essential for Plant Autophagy. Plant Cell 16: 2967–2983.

Yoshimoto K. 2012. Beginning to understand autophagy, an intracellular self-degradation system in plants. Plant and Cell Physiology. 53: 1355–1365.

Zhuang X, Chung KP, Cui Y, Lin W, Gao C, Kang BH, Jiang L. 2017. ATG9 regulates autophagosome progression from the endoplasmic reticulum in Arabidopsis. Proceedings of National Academy of Science of the United States of America 114: E426–E435.

