## Supplemental file for "Autophagy is induced during plant grafting for wound healing"

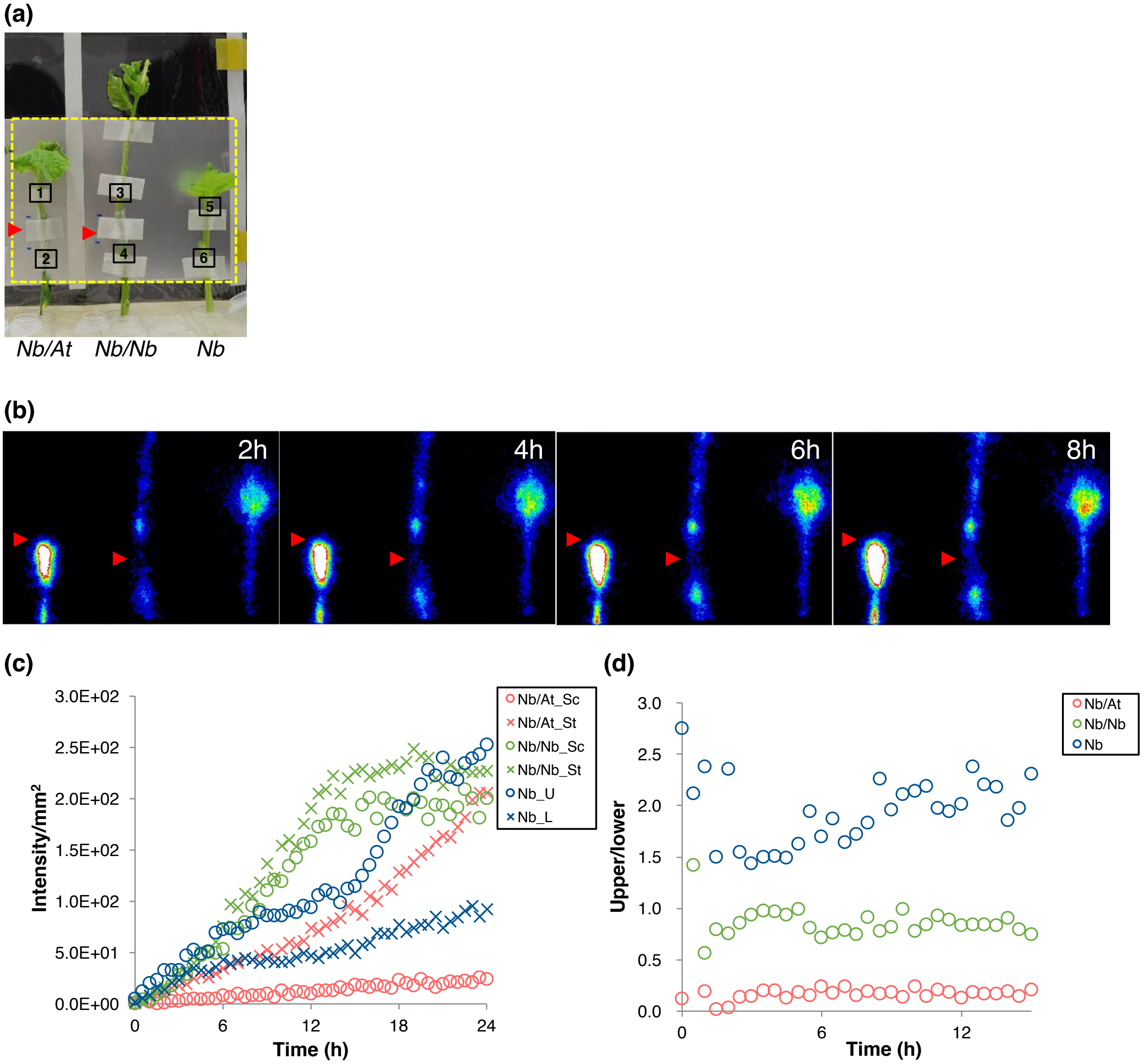

**Fig S1** Radio isotope transport experiments in the intact *Nb* plants, *Nb*/*Nb* homografts and *Nb*/*At* heterografts. (a) The imaging apparatus for the Pi labeled with $^{32}P$ flow detection. Rectangle indicates the imaging area shown in (b). Arrowheads indicate the position of grafting. Boxes with numbers indicate the measuring points for (c): 1, *Nb*/*At*_Scion (Sc); 2, *Nb*/*At*_Stock (St); 3, *Nb*/*Nb*_Sc; 4, *Nb*/*Nb*_St; 5, intact *Nb*_Upper (U); 6, intact *Nb*_Lower (L). (b) The time course imaging of Pi flow. Arrowheads indicate the positions of grafting. (c) The time course plots for radio isotope signal intensity per unit area. (d) The time course plots for the values of *Nb*/*At*_Sc (1)/*Nb*/*At*_St (2), *Nb*/*Nb*_Sc (3)/*Nb*/*Nb*_St (4), and *Nb*_U (5)/*Nb*_L (6) in (c).

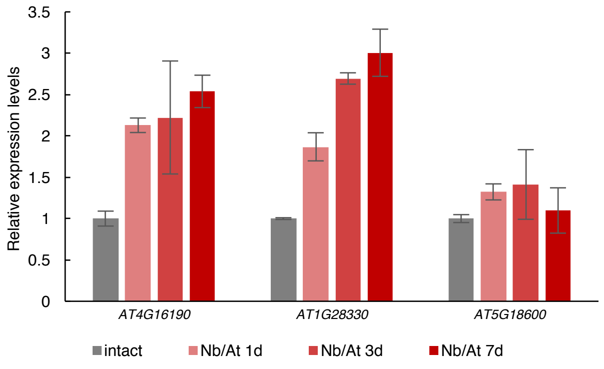

**Fig. S2** Relative expression of the genes related to nutrient deficiency in the *At* stock part of *Nb*/*At* heterografts. The expression levels of *At* genes were established by real-time PCR analysis in the intact *At* and *Nb*/*At* heterograft samples. Expression levels were normalized by the levels of *EF1-a* and adjusted to be relative to the the intact samples. RNA was extracted from the stem tissues of 10 intact or grafted plants for each sample fractions. Mean ± s.d. from 3 independent experiments.

**Table S1** The sequences of the PCR primers used in this study

| | Primer name | sequence |
|---|---|---|
| for Quantitative reverse-transcription PCR | | |
| | Niben101Scf12671g00007_1271-1288_FWD_1 | 5'-TCGATCATGGGCCTGGCA-3' |
| | Niben101Scf12671g00007_1365-1384_REV_1 | 5'-CCTCCTCCTCCTGTCAACCC-3' |
| | Niben101Scf05415g00003_156-173_FWD_1 | 5'-ACGCGCCTCTGCTTAGCT-3' |
| | Niben101Scf05415g00003_309-327_REV_1 | 5'-GGGTTTATGTGGCCTTGGC-3' |
| | Niben101Scf00390g02001_87-104_FWD_1 | 5'-ACAAATTCCGCCGCCCAC-3' |
| | Niben101Scf00390g02001_237-254_REV_1 | 5'-TGGTGTCGTGACGGTGGT-3' |
| | NbATG5-F | 5'-GAAGCTTATCTCCGAATCTCGTC-3' |
| | NbATG5-R | 5'-TGAGTGCTTTCCGGTGATTTA-3' |
| | NbACT1-F | 5'-GGCCAATCGAGAAAAGATGAC-3' |
| | NbACT1-R | 5'-AACTGTGTGGCTGACACCATC-3' |
| | AT1G28330_93-112_FWD_1 | 5'-CCTTGGCCGCCTCCGTAAGA-3' |
| | AT1G28330_178-197_REV_1 | 5'-GCCGGCATGGTCAACGACCT-3' |
| | AT1G76520_828-849_FWD_1 | 5'-CCAAAAGGCTGGGAGGTGGGAA-3' |
| | AT1G76520_982-1001_REV_1 | 5'-ACCCGGAGAGGAGCTTCCGT-3' |
| | AT5G18600_347-366_FWD_1 | 5'-TGTTGCGGCTCGGGTGTAGC-3' |
| | AT5G18600_466-485_REV_1 | 5'-ACCCACAATGCACCAGCCCG-3' |
| | AtEF1a_qPCR_F1 | 5'-TACTGGTACCTCCCAGGCTGA-3' |
| | AtEF1a_qPCR_R1 | 5'-ACACCAAGGGTGAAAGCAAGA-3' |
| for VIGS construction of NbATG5 | | |
| | NbATG5_VIGS_F | 5'-CTTAAAATCTCAGCTTCACCA-3' |
| | NbATG5_VIGS_R | 5'-AGGTTACCTGCCTCTTTTAGGAACA-3' |
| for RT-PCR reaction in VIGS experiments | | |
| | VIGS_RT-PCR_F | 5'-ATTCAGATCGTCGTCAGTGC-3' |
| | VIGS_RT-PCR_R | 5'-AGCAATACTGCCAACTCAGC-3' |

**Table S2** Comparisons of the success rates of At wild type and atg2 mutant self-grafts

| | #1 | | #2 | | #3 | | #4 | | #5 | |
|---|---|---|---|---|---|---|---|---|---|---|
| | WT | *atg2* | WT | *atg2* | WT | *atg2* | WT | *atg2* | WT | *atg2* |
| Grafted plants | 38 | 48 | 31 | 38 | 11 | 47 | 35 | 70 | 36 | 31 |
| Succeeded grafts | 23 | 23 | 16 | 19 | 5 | 28 | 16 | 40 | 19 | 22 |
| Success rate (%) | 60.5 | 47.9 | 51.6 | 50.0 | 45.5 | 59.6 | 45.7 | 57.1 | 52.8 | 71.0 |
| $\chi^2$ value | 0.730 | | 0.018 | | 0.725 | | 1.224 | | 2.321 | |
| *P* value | 0.393 | | 0.894 | | 0.395 | | 0.268 | | 0.128 | |

The success rates of wild type and *atg2* mutant self-grafts in five independent experiments were shown. From the top low, the number of grafted plants, the number of succeeded grafts and success rate of grafting (%) are shown. Using the chi-square test, it was tested whether there was a significant difference in the survival rates between wild type and *atg2* mutant self-grafts. $\chi^2$ and *P* values for each experiment are shown.
